# Comprehensive antibody profiling of mRNA vaccination in children

**DOI:** 10.1101/2021.10.07.463592

**Authors:** Yannic C Bartsch, Kerri J St Denis, Paulina Kaplonek, Jaewon Kang, Evan C. Lam, Madeleine D Burns, Eva J Farkas, Jameson P Davis, Brittany P Boribong, Andrea G Edlow, Alessio Fasano, Wayne Shreffler, Dace Zavadska, Marina Johnson, David Goldblatt, Alejandro B Balazs, Lael M Yonker, Galit Alter

## Abstract

While children have been largely spared from COVID-19 disease, the emergence of viral variants of concern (VOC) with increased transmissibility, combined with fluctuating mask mandates and school re-openings have led to increased infections and disease among children. Thus, there is an urgent need to roll out COVID-19 vaccines to children of all ages. However, whether children respond equivalently to adults to mRNA vaccines and whether dosing will elicit optimal immunity remains unclear. Given the recent announcement of incomplete immunity induced by the pediatric dose of the BNT162b2 vaccine in young children, here we aimed to deeply profile and compare the vaccine-induced humoral immune response in 6-11 year old children receiving the pediatric (50μg) or adult (100μg) dose of the mRNA-1273 vaccine compared to adults and naturally infected children or children that experienced multi inflammatory syndrome in children (MIS-C) for the first time. Children elicited an IgG dominant vaccine induced immune response, surpassing adults at a matched 100μg dose, but more variable immunity at a 50μg dose. Irrespective of titer, children generated antibodies with enhanced Fc-receptor binding capacity. Moreover, like adults, children generated cross-VOC humoral immunity, marked by a decline of omicron receptor binding domain-binding, but robustly preserved omicron Spike-receptor binding, with robustly preserved Fc-receptor binding capabilities, in a dose dependent manner. These data indicate that while both 50μg and 100μg of mRNA vaccination in children elicits robust cross-VOC antibody responses, 100ug of mRNA in children results in highly preserved omicron-specific functional humoral immunity.

**One-Sentence Summary:** mRNA vaccination elicits robust humoral immune responses to SARS-CoV-2 in children 6-11 years of age.

## Introduction

The burden of respiratory infections is often higher in young children with a developing untrained immune system(*1*). However, lower rates of disease were noted early in the COVID-19 pandemic among children, who largely experienced asymptomatic or pauci-symptomatic SARS-CoV-2 infections(*2*). However, with the rise of highly infectious variants of concern (VOCs), like the novel omicron VOC, increasing infection rates and hospitalizations have been observed globally(*3, 4*). Linked to the unpredictable incidence of multisystem inflammatory syndrome in children (MIS-C) and the clear contribution children make to population level spread, the need for vaccines for children is evident (*5, 6*). However, whether newly emerging COVID-19 vaccine platforms, approved for teenager/adult use, elicit immunity in children is not well understood.

Epidemiologic data clearly highlight vulnerabilities in the pediatric immune system, with increased rates of respiratory, enteric, and parasitic infections disproportionately causing disease in children in the first decade of life(*7, 8*). In fact, vaccine-induced immune responses often differ across children and adults(*9*). However, whether these vulnerabilities to infection and poor response to protein-based vaccination will translate to newer vaccine platforms, like mRNA vaccine platforms, remains unclear. Moreover, emerging data suggests that dosing may not be straight forward for mRNA vaccines (*10, 11*), due to reduced immunogenicity in young children, requiring deeper immunologic insights to guide rational pediatric vaccine design.

To begin to define the humoral mRNA vaccine responses in children, we comprehensively profiled vaccine-induced immune responses in children (6-11 years) who received the pediatric (50μg) or adult (100μg) dose of the mRNA-1273 vaccine regimen, respectively. We observed 100% vaccine response rates prior to the second vaccine dose in children that received the 100μg vaccine dose. While immune profiles in the low (50μg) dose were more similar to adults (who received the adults recommended 100μg dose), children receiving the adult (100μg) dose generated disproportionately higher IgG biased vaccine responses following the second vaccine dose, with enhanced Fc-effector profiles. Moreover, both pediatric and adult doses elicited broad cross-variant isotype and Fc-receptor binding antibodies, however, while all groups experienced a significant loss of omicron-receptor binding domain (RBD) reactivity, omicron Spike-specific immunity was largely preserved and 100μg immunized children exhibited the highest cross-reactivity. Collectively, these data point to robust, but dose dependent, functional humoral pediatric immune signatures induced in children following mRNA-1273 vaccination.

## Results

In the wake of fluctuating mask mandates, school re-openings, and the rapid spread of the highly infectious SARS-CoV-2 delta and omicron variants, a surge of SARS-CoV-2 infections in children has been observed (*5*). Increasing numbers of children with severe COVID-19 or life-threatening Multisystem Inflammatory Syndrome in Children (MIS-C), plus our evolving appreciation of children in the spread of the pandemic, there is an urgent need to roll out vaccines across all ages. With the rapid roll out of mRNA vaccines, it remains unclear whether children will generate sufficiently robust immunity following mRNA vaccination. Here we deeply characterized the immune response induced by the Moderna mRNA-1273 vaccine in children that received an adult dose (100μg) of mRNA-1273 (n=12; median age= 9 years range: 7 – 11 years; 42% female), matching the recommendations for adults, or a pediatric (50μg) dose of mRNA-1273 (n=12; median age= 8 years range: 5 – 11 years; 50% female) at days 0 and 28, respectively. Plasma samples were collected before vaccination (V0), approximately four weeks after prime (V1) and four weeks after second (V2) immunization.

### mRNA vaccines induce robust SARS-CoV-2 spike binding and neutralizing titers in children

To begin to investigate the vaccine-induced humoral response, we profiled SARS-CoV-2 Spike (S) specific antibody titers. At V1, we observed seroconversion (marked by an increase in Spike specific IgM, IgA1 or IgG1 binding compared to V0) in 100 % of children receiving the 50μg (n=9/9) or 100μg (n=12/12) dose of mRNA-1273. Both S-specific IgA1 and IgG1 increased with the second dose, while S-specific IgM responses declined slightly, marking efficient class switching. After the second dose we observed significantly elevated S-specific IgA1 levels in adults compared to children (p-value: <0.001) (**Figure 1A**). In contrast, children in the 100μg dose group elicited higher IgG1 titers after the first and second dose of the vaccine compared to children in the 50μg dose group (p-value: 0.004), as well as compared to vaccinated adults (p-value: 0.03). Likewise, we observed a trend towards higher neutralizing antibody titers in the pediatric 100μg dose group followed by adults and 50μg vaccinated children (**Figure 1B**). Univariate comparison of V2 levels of the 100μg adults to 100μg and 50μg children highlighted isotype selection differences across children and adults, but minimal overall differences in antibody binding titers and neutralization across the 50 and 100μg doses in children (**Figure 1C**). Furthermore, vaccine-induced binding and neutralization titers in the 100μg pediatric dose group were higher compared to levels observed in naturally exposed convalescent children or MIS-C. Importantly, while these differences may be related to exposure to different variants, we observed superior vaccine induced binding and neutralization to all variants, highlighting the critical importance of SARS-CoV-2 vaccination in promoting broader VOC immunity in children compared to infection (**Supplemental Figure 1 and 2**). Taken together, these data show that mRNA vaccination can elicit strong but dose-dependent anti-SARS-CoV-2 binding and neutralizing titers in children superior to natural infection that are accompanied by some age-dependent shifts in isotype-antibody profiles.

**Figure 1.**
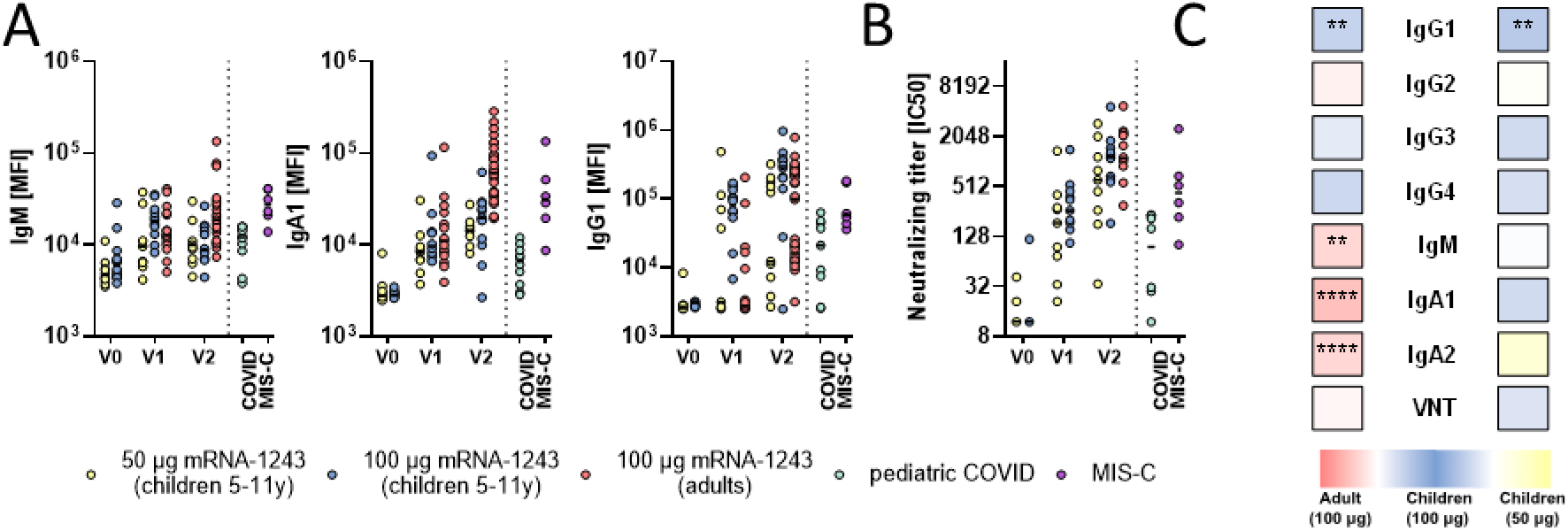
mRNA-1273 vaccination induces robust binding and neutralizing titers in children. A) Relative SARS-CoV-2 spike (Wuhan) specific IgM, IgA1 and IgG1 binding levels were determined by Luminex in children (6-11 years) receiving 50μg or 100μg mRNA1273 before (V0_50_: n=12; V0_100_: n=12), after the first (V1_50_: n=9; V1_100_: n=12) or after the second (V1_50_: n=11; V2_100_: n=12) dose or in adults receiving two 100μg doses (V1: n=19; V2: n=33) as well as in convalescent pediatric COVID (n=9) or MIS-C (n=6). B) The dot plots show the inverse 50 % virus neutralizing titers in children (6-11 years) receiving 50μg or 100μg mRNA1273 before (V0_50_: n=12; V0_100_: n=12), after the first (V1_50_: n=9; V1_100_: n=12) or after the second (V1_50_: n=9; V2_100_: n=11) dose or in adults receiving two 100μg doses (V2: n=14) as well as in convalescent pediatric COVID (n=9) or MIS-C (n=6). C) Heatmap strips summarize univariate comparison at the V2 timepoint of 100μg dose vaccinated children to adults (left panel) or to 50μg dose vaccinated children (right panel). Color of the tiles indicate whether antibody binding titer were upregulated in the respective group: 100μg vaccinated children (blue shades), adults (red shades), or 50μg vaccinated children (yellow shades). A Wilcoxon-rank test was used to test for statistical significance and asterisks indicate statistically significant differences of the respective feature after Benjamini-Hochberg correction for multiple testing (*:p<0.05; **:p<0.01;***:p<0.001).

### mRNA vaccination induces highly potent Spike-specific Fc effector functions in children

In addition to binding and neutralization, protection against severe adult COVID-19 has been linked to the ability of antibodies to leverage additional antiviral functions via Fc-receptors to fight infection, referred to as antibody effector functions(*12–14*). Specifically, opsonophagocytic pathogen clearance is key to protection against several bacterial pathogens and cytotoxic antibody functions have been linked to protection against viruses(*15, 16*). Thus, we profiled the relative ability of vaccine-induced immune responses to bind to human Fc-receptors (FcγR2a, FcγR2b, FcγR3a, FcγR3b, and FcαR) as well as their ability to elicit antibody-dependent complement deposition (ADCD), antibody dependent neutrophil phagocytosis (ADNP), antibody dependent monocyte phagocytosis (ADCP), or activation antibody dependent NK cell activation (ADNKA). Children in both dose groups elicited Spike (S)-specific IgG antibodies that bound robustly to all Fc-receptors following the first dose, markedly greater than responses observed in vaccinated adults and in natural COVID-19 infection or MIS-C (**Figure 2A**). Moreover, these responses expanded further after the second immunization with significantly elevated antibody-Fc-receptor binding in the 100μg pediatric dose group compared to the 50μg dose, with significant higher FcγR3a and slightly higher FcγR2a, FcγR2b and FcγR3b binding compared to the 100μg adult group (**Figure 2B**). Interestingly, this increased Fc-receptor binding was not directly related to overall changes in Spike-specific IgG subclass selection (**Supplemental Figure 1**), pointing to alternate mechanisms for augmented humoral immune function in children, potentially linked to pediatric selection of more potent Fc-glycosylation profiles(*17*). In contrast, compared to adults, children induced lower levels of IgA antibodies that exhibited, as expected, lower interactions with the IgA-Fc-receptor, FcαR, compared to adults (**Figure 2B and Supplemental Figure 2**)(*18, 19*).

**Figure 2.**
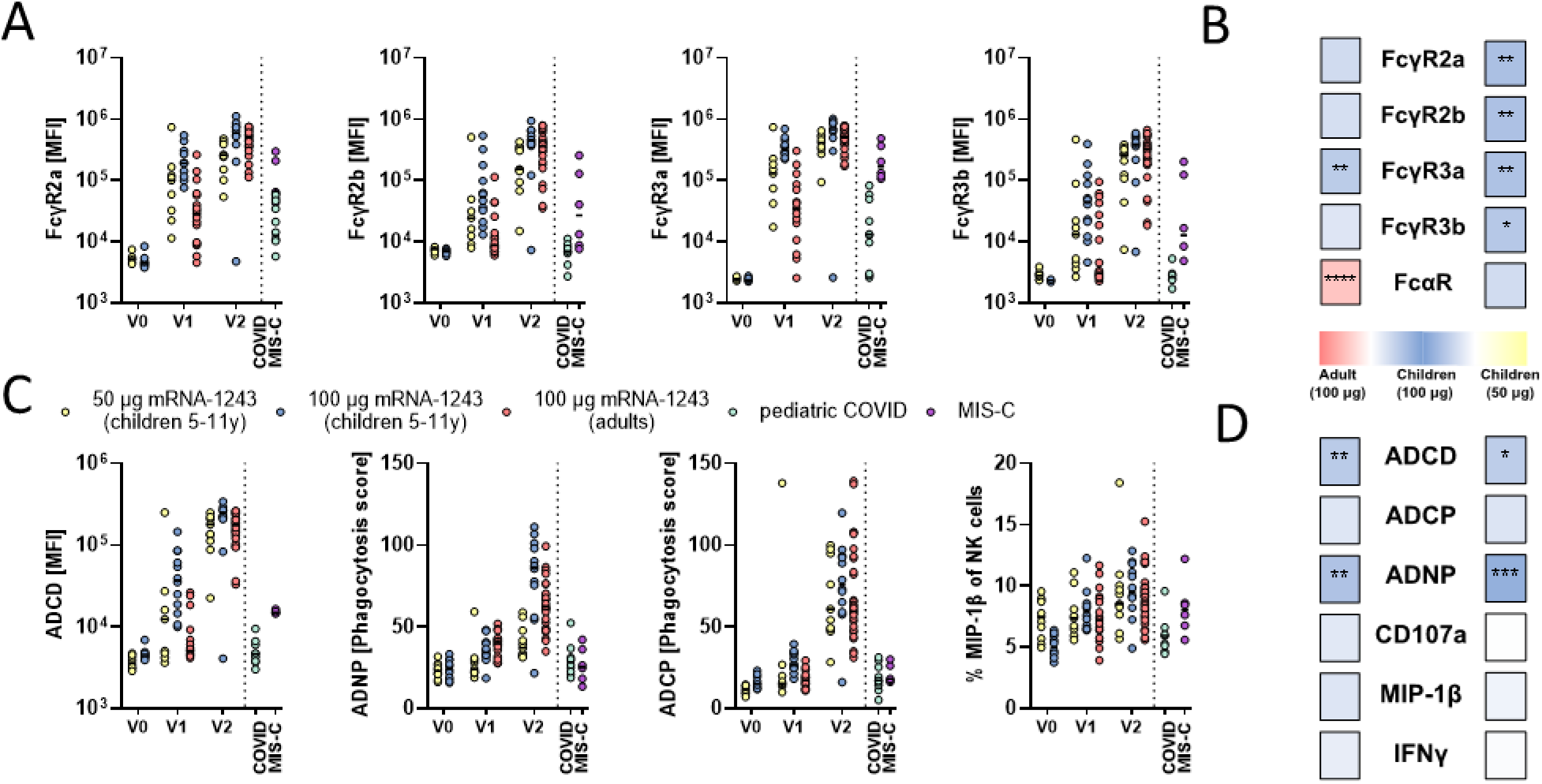
mRNA-1273 vaccination induces higher FcγR binding and phagocytic activity in children. A) Binding of SARS-CoV-2 specific antibodies to FcγR2a, 2b, 3a, and 3b was determined by Luminex in children (6-11 years) receiving 50μg or 100μg mRNA1273 before (V0_50_: n=12; V0_100_: n=12), after the first (V1_50_: n=9; V1_100_: n=12) or after the second (V1_50_: n=11; V2_100_: n=12) dose or in adults receiving two 100μg doses (V1: n=19; V2: n=33) as well as in convalescent pediatric COVID (n=9) or MIS-C (n=6). C) Heatmap strips summarize univariate comparison of Fc receptor binding at the V2 timepoint of 100μg dose vaccinated children to adults (left panel) or to 50μg dose vaccinated children (right panel). Color of the tiles indicate whether antibody binding titer were upregulated in the respective group: 100μg vaccinated children (blue shades), adults (red shades), or 50μg vaccinated children (yellow shades). C) The ability of SARS-CoV-2 S specific antibody Fc to induce antibody-dependent-complement-deposition (ADCD), neutrophil-phagocytosis (ADNP), cellular-THP1 monocyte-phagocytosis (ADCP), or activation of NK cells marked by expression of MIP-1β was analyzed. D) Heatmap strips summarize univariate comparison of Fc effector functions at the V2 timepoint of 100μg dose vaccinated children to adults (left panel) or to 50μg dose vaccinated children (right panel) as in B). A Wilcoxon-rank test was used to test for statistical significance in C) and D) and asterisks indicate statistically significant differences of the respective feature after Benjamini-Hochberg correction for multiple testing (*:p<0.05; **:p<0.01;***:p<0.001).

To next determine whether these distinct pediatric Fcγ-receptor binding profiles translated to more functional Spike-specific humoral immune responses, we examined vaccine-induced Fc-effector functions (**Figure 2C**). Interestingly, low levels of ADCD, ADNP and ACDP were observed after primary immunization, but were notably augmented by the second immunization across the groups, resulting in significantly increased ADCD and ADNP in the 100μg vaccinated children compared to adults or the 50μg pediatric dose (**Figure 2C**). In contrast, NK cell functions (as measured by MIP-1b expression) were induced to equal levels across all groups. Overall, high dose mRNA-1273 induced higher levels of ADCD, ADNP and ADCP recruiting antibodies in children compared to adults following the 100μg vaccine regimen (**Figure 2D**). These data point to enhanced functional antibody responsiveness in children at a matched 100μg dose compared to adults, and a solid functional response at the optimized 50μg dose endowing children with a robust capacity to recruit immune function at half the adult dose, all higher than levels observed following natural infection or MIS-C.

### Selective expansion of opsonophagocytic antibodies with mRNA vaccination in children

To gain a more granular sense of the differences in immune responses across children and adults at the same matched dose or across children receiving the 50 and 100μg doses, we next utilized a machine learning approach to probe the humoral immune features that differed most across these groups. As few as six of the overall features analyzed across all plasma samples, were sufficient to completely resolve vaccine induced immune responses induced by the 100μg dose across children and adults (**Figure 3A+B**). Specifically, vaccine-induced S-specific IgG1, FcγR3a binding, ADNP and ADCD were all enriched selectively in children, whereas Spike-specific IgM and IgA1 titers were enriched in adults (**Figure 3A**), highlighting distinct isotype selection in adults, and the generation of more functional antibodies in children. Conversely, comparison of 50 and 100μg doses in children was achieved using only two of all antibody features analyzed for each plasma. These features Spike-specific ADNP and IgG4 levels, both of which were enriched in the immune profiles in children that received the 100μg dose of the vaccine (**Figure 3C+D**). Additionally, in contrast to natural infection, vaccination induced higher titers and functions to SARS-CoV-2 when comparing vaccinated children to those that were previously infected (**Figure 2 and Supplemental Figure 3**). Collectively, these data point to slight shifts in isotype selection between adults and children, but the potential for children to raise more functional antibodies, that match those of adults at a half-dose.

**Figure 3.**
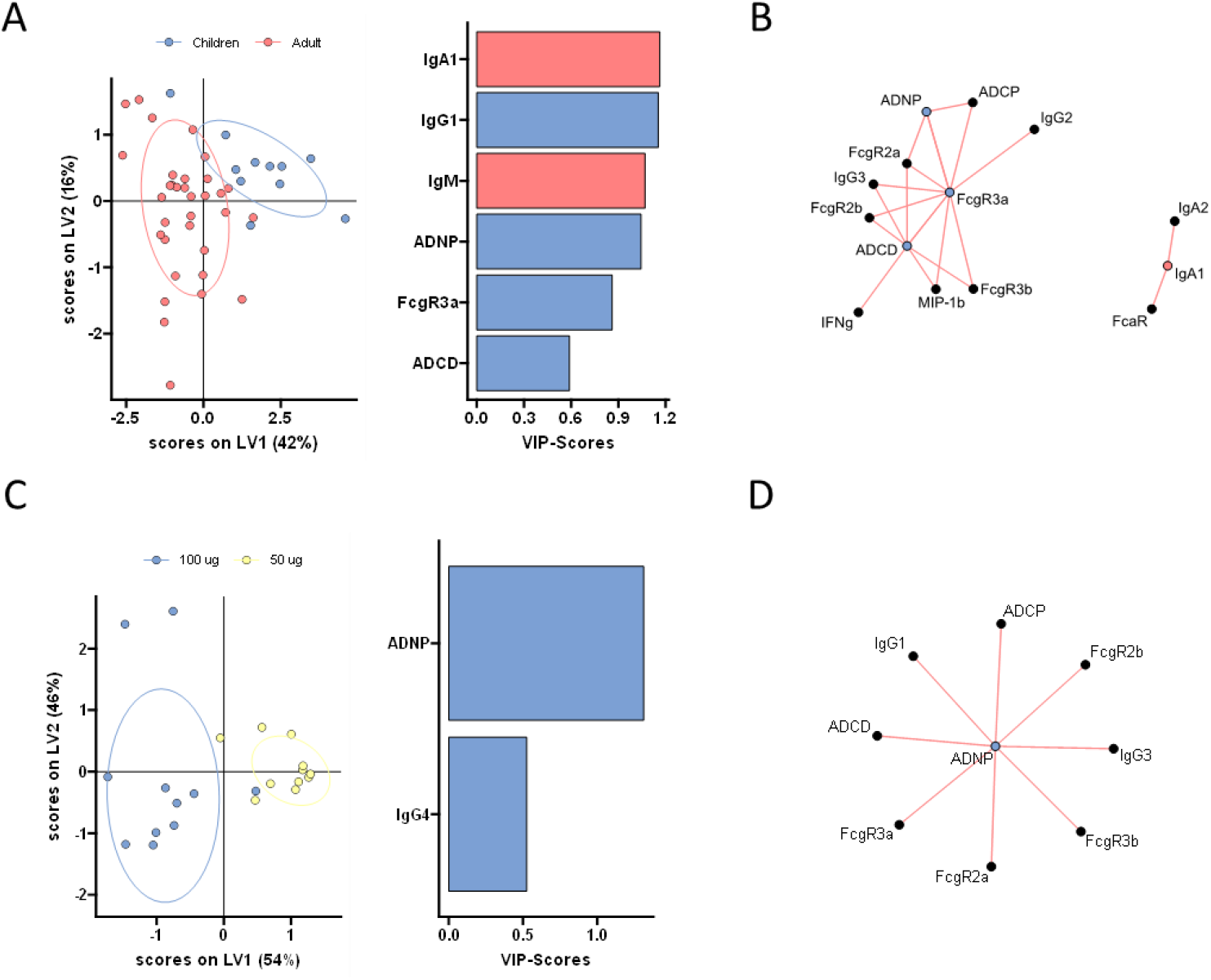
Distinct humoral profiles distinguish between adult and pediatric vaccine responses. A) A machine learning model was built using a minimal set of LASSO selected SARS-CoV-2 S specific features at V2 (left panel) to discriminate between vaccine responses in adult (red) and 100 μg vaccinated children (purple) in a PLS-DA analysis (right panel). B) The co-correlation network illustrates all LASSO-selected features. Nodes of selected features are colored whether they were enriched in children (purple) or adults (red). Lines indicate significant (p<0.05) spearman correlations with |r|>0.7 of connected features (only positive correlations with r>0 were observed). C) PLS-DA model of LASSO selected features at V2 (left panel) to discriminate between vaccine responses in 100 μg (purple) and 50 μg (yellow) vaccinated children. D) The co-correlation network as in B) illustrates all LASSO-selected features. Nodes of selected features are colored whether they were enriched in 100 μg vaccinated children (purple).

### mRNA vaccination in children raises robust responses against SARS-CoV-2 variants of concern

Real-world efficacy suggests that mRNA vaccines confer robust protection against severe disease/death against the original (wildtype; wt) SARS-CoV-2 strain (Wuhan), at levels greater than 90% (*20*). This level of efficacy appears to be sustained against evolving variants of concern (VOCs), including the alpha and delta variants, although lower levels of protection have been observed against the beta variant (*21, 22*). While previous VOCs were marked by single or few amino acid substitutions, the novel omicron variant has 29 mutations in the Spike protein, resulting in enhanced transmissibility, and a concomitant loss of neutralizing titers (*23, 24*). Yet, despite the striking increase in omicron transmissibility, a similar increase in severe disease and death has not been observed, suggesting that alternate vaccine induced immune responses may continue to afford protection against severe disease and death. To explore whether mRNA vaccination in children results in the generation of vaccines with differential VOC recognition capabilities (*25*). We observed a progressive loss of IgM, IgA, and IgG binding to VOC RBDs across both pediatric groups and adults, with more variable cross-VOC IgG responses among 50μg immunized children, but a consistent and significant loss of binding to the omicron RBD across all 3 groups (Figure 4A). Conversely, Spike-specific responses were more resilient across most VOCs, across the 3 groups, except for omicron-Spike-specific responses that were significantly lower across IgM and IgA response across the groups (Figure 4B). Yet, IgG responses showed 3 different patterns: 1) 50μg immunized children experienced heterogeneous responses across VOCs, marked by some of the lowest omicron-responses, 2) adults exhibited more stable Spike VOC-IgG binding levels, but experienced a significant reduction in omicron-Spike reactivity, and 3) 100μg immunized children exhibited negligible reduction in Spike-specific recognition across VOCs, including to the omicron Spike. Furthermore, Fc-receptor binding capability was largely preserved across RBD VOCs, except for omicron RBD-binding which was significantly lower across all 3 groups (Figure 4C). However, Spike-VOC binding differed across the 3 groups. While wildtype, alpha, beta, and delta VOC Fc-receptor binding profiles were highly preserved across all 3 groups, omicron-Spike Fc-receptor binding was most significantly lost in 50μg immunized children (Figure 4D). Adults exhibited an intermediate loss of omicron-Spike Fc-receptor binding, with a more pronounced preservation of the opsinophagocytic FcγR2a and cytotoxic FcγR3a binding. Conversely, 100μg immunized children exhibited a negligible loss of Fc-receptor binding to the omicron Spike, pointing to the generation of highly resilient antibodies in children at the adult dose that continue to bind to the omicron Spike despite the significant loss of RBD binding. Whether children generate broader or more flexible humoral immune response at the 100μg dose, enabling them to preserve immunity to VOCs remains unclear, but the data point to dose and age dependent effects of antibody-mediated cross-VOC recognition. Whether differences in neutralization and Fc-function lead to differences in disease-breakthrough across the ages remains unclear but provides some additional immunological insights that may continue to explain the epidemiologic differences in disease severity globally in the setting of emerging VOCs.

**Figure 4.**
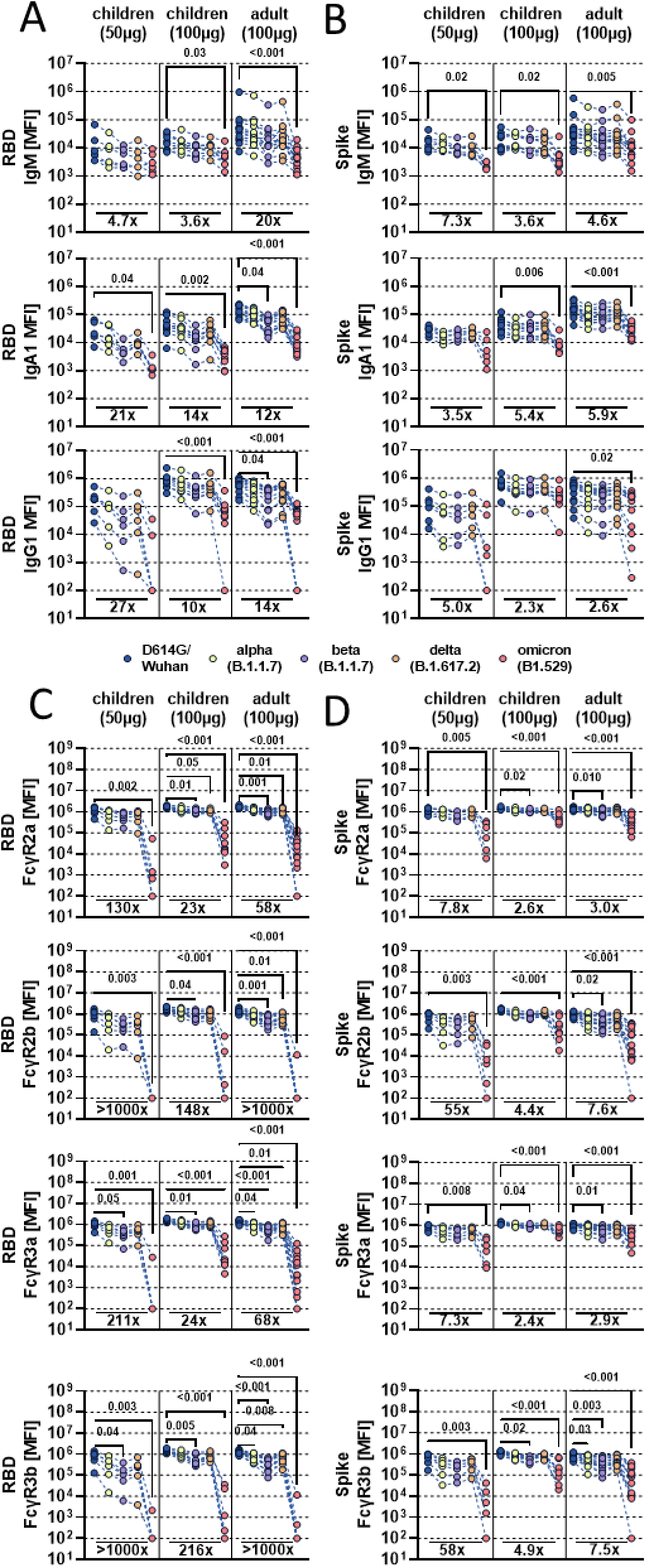
mRNA-1273 vaccination elicits humoral responses to SARS-CoV-2 variants of concern. A-B) The line graphs show the vaccine induced IgM, IgA1 and IgG1 recognition to D614G (WT; blue), alpha (B1.117; yellow), beta (B1.351; purple), delta (B.1.617.2; orange), and omicron (B1.529; red) variants of concern receptor binding domains (A) or full Spike (B) for children (n_50μg_=6, n_100μg_=9) or adults (n=14) at V2, where each individuals’ response is linked across VOC antigens. C-D) The line graphs show the FcγR (FcγR2a, FcγR2b, FcγR3a, FcγR3b) binding profiles of vaccine induced antibodies to RBD or Spike VOC antigens across children (n_50μg_=6, n_100μg_=9) or adults (n=12) at V2, where each individuals’ response is linked across VOC antigens. Background corrected data is shown and negative values were set to 100 for graphing purposes in A-D. A Kruskal-Wallis test with a Benjamini-Hochberg post-test correction for multiple comparisons was used to test for statistical differences between wildtype and VOC titers within groups. P-values for significantly different features are shown above and fold change reduction of omicron titer compared to wildtype below each dataset.

## Discussion

mRNA vaccine platforms responded rapidly to SARS-CoV-2 threat, demonstrating robust levels of efficacy in adults(*26, 27*). However, despite the successes of SARS-CoV-2 vaccines, the global roll out has begun to highlight key vulnerable populations, and strategic gaps, that may limit the impact of vaccination globally. Although children generally experience mild symptoms, they can harbor robust, high levels of SARS-CoV-2 replication, thereby contributing significantly to viral *spread*(*28–30*). Furthermore, increasing numbers of children are suffering from severe COVID-19 with over 28,000 hospitalizations and over 700 deaths in the US alone as of December 2021 (*31*). However, because children have a more naïve immune system that evolves with age it was uncertain how the mRNA vaccine platforms would impact immunogenicity in young children. Additionally, recent results suggest that dose adjustments for very young children, due to safety and tolerance concerns, have introduced additional variation in immunogenicity, resulting in poor immunogenicity in children under 5 years, who received a lower dose than the adult recommended 30μg dose (*11*). Thus, in the absence of empirical data, optimal dosing is uncertain. Thus, here we aimed to dive deeply in defining humoral profile differences across doses and across children and adults. Similar to results with the Pfizer and Moderna mRNA vaccine trials in teenagers (*32, 33*), here we found that the Moderna mRNA vaccine was highly immunogenic in 6-11 year old children, generating a humoral response superior to that seen following viral exposure. However, granular vaccine-induced humoral profiling identified significant differences in adult and pediatric vaccine responses, marked by a selective induction of highly functional IgG responses, with fewer IgA and IgM responses compared to adults. At a matched 100μg dose, children mounted more robust opsonophagocytic functions, and Fcγ-receptor binding compared to adults. Moreover, at half the adult dose, children mounted equal, albeit more variable, responses compared to adults.

Virus neutralization represents a key surrogate marker of vaccine protection against COVID-19. Yet despite the loss of neutralization against several emerging variants of concern (VOCs), mRNA vaccines continue to provide protection against severe disease and death(*21, 34*). Interestingly, opsonophagocytic functions of antibodies, rather than neutralization alone, have been linked to survival of COVID-19 following natural infection(*12*) and are associated with protection from infection in animal models(*35, 36*). Here, when immunized with the adult dose, children induced comparable neutralization but exhibited a preferential expansion of opsonophagocytic functions compared to adults. The enhanced opsonophagocytic function was not linked to differential subclass or isotype selection, suggesting that children may induce more functional antibodies via alternate changes to the humoral immune response, including potential differences in post-translational IgG modification that may lead to more flexible, highly functional responses, representing an evolutionary adaptation enabling children to react more flexibly to infections(*37*).

The adaptive immune response matures during the first decade of life(*38*). Several lines of evidence suggest that the more naïve immune response in children may allow the immune system to adapt and evolve more easily to new pathogens, poised to generate broader immunity to new viruses(*39, 40*). Moreover, throughout life, our naïve clonal repertoire or immune cells shift in response to the sequence of pathogens and vaccines we are exposed to. Thus, naïve children may have a less “biased” repertoire, enabling the generation of immunity to a broader range of pathogens(*41*). Along these lines, we observed robust induction of immunity against most VOCs, with the exception of omicron. However, IgG and Fc-receptor binding profiles were highly similar among children and adults, although 100μg immunized children induced IgG that were more resilient against VOCs, exhibiting robust recognition of the omicron Spike, whereas both 50μg immunized children and 100μg immunized adults both lost partial recognition of the omicron Spike. These data suggest that higher pediatric dosing can result in more flexible humoral immunity in children against highly divergent VOCs, superior to those induced in adults at a matched dose. Thus, children may generate more functional Fc-effector functions, that while not neutralizing, may be poised for rapid elimination of the pathogen upon transmission, providing a highly effective means to prevent COVID-19.

With the increasing spread of SARS-CoV-2 omicron among younger populations (*42*), increasing incidence of COVID-19 among the pediatric population, the rare incidence of MIS-C, and the recent appreciation for long-COVID in children, the need to determine whether SARS-CoV-2 vaccines can elicit functional immune responses will be key to protect children(*3, 5, 28, 43, 44*). Comparable to previous observations in adults(*10*), the mRNA-1273 vaccine induced robust binding titers, neutralization, and Fc effector function in vaccinated children, in a dose dependent manner, compared to children diagnosed with COVID-19 or MIS-C, pointing to the importance of vaccination to robustly bolster immunity to SARS-CoV-2 and emerging VOCs to provide broader and more potent immunity to SARS-CoV-2. Whether these responses will wane differentially across doses, whether they will be more protective against particular VOCs, and whether children will require boosting remains unclear, however, these findings support vaccination of children with mRNA-1273 as a safe and effective strategy to protect children against COVID-19, MIS-C, and Long-COVID.

## Contributions

Y.C.B. and J.K. performed the serological experiments. K.J.D, E.C.L and A.B.B. performed the neutralization assay. Y.C.B., L.M.Y. and G.A. analyzed and interpreted the data. M.D.B., E.J.F., J.P.D., B.P.B., A.G.F., A.F., W.S., D.Z., M.J., D.G., and L.M.Y. supervised and managed the sample collection. G.A. supervised the project. Y.C.B., L.M.Y. and G.A. drafted the manuscript. All authors critically reviewed the manuscript.

## Acknowledgment

We thank Nancy Zimmerman, Mark and Lisa Schwartz, an anonymous donor (financial support), Terry and Susan Ragon, and the SAMANA Kay MGH Research Scholars award for their support. We acknowledge support from the Ragon Institute of MGH, MIT, and Harvard, the Massachusetts Consortium on Pathogen Readiness (MassCPR), the NIH (3R37AI080289-11S1, R01AI146785, U19AI42790-01, U19AI135995-02, U19AI42790-01, 1U01CA260476 – 01, CIVIC75N93019C00052, 5K08HL143183), the Gates Foundation Global Health Vaccine Accelerator Platform funding (OPP1146996 and INV-001650), and the Musk Foundation.

## Methods

### Cohort

Pediatric vaccinee samples were obtained from children who were vaccinated with two doses 100 μg mRNA-1273 at MGH as participants in Part 1 (open label) of a Phase2/3 clinical trial (ClinicalTrials.gov Identifier: NCT04796896). Additionally, we included samples from eight children who presented with acute PCR confirmed COVID-19 (7-19 years) or six children with MIS-C (3-22 years) at our hospital. Additionally, samples from 14 adults who received two doses mRNA-1273 as part of a phase 1 clinical trial were used as controls (ClinicalTrials.gov Identifier: NCT04283461). All pediatric participants provided informed assent and their legal guardian provided informed consent prior to participation in the MGH Pediatric COVID-19 Biorepository. Blood samples and symptom report via an IRB-approved symptom questionnaire were collected prior to vaccination, one month after the first vaccination and one month after the second vaccination. This study was overseen and approved by the MassGeneral Institutional Review Board (IRB #2020P00955).

### Antigens and biotinylation

All antigens were biotinylated using an NHS-Sulfo-LC-LC kit according to the manufacturer’s instruction (Thermo Fisher, MA, USA) if required by the assay and excessive biotin was removed by size exclusion chromatography using Zeba-Spin desalting columns (7kDa cutoff, Thermo Fisher).

### Antibody isotype and Fc receptor binding

Antigen-specific antibody isotype and subclass titers and Fc receptor binding profiles were analyzed with a custom multiplex Luminex assay as described previously (*45*). In brief, antigens were coupled directly to Luminex microspheres (Luminex Corp, TX, USA). Coupled beads were incubated with diluted plasma samples washed, and Ig isotypes or subclasses with a 1:100 diluted PE-conjugated secondary antibody for IgG1 (clone: HP6001), IgG2 (clone: 31-7-4), IgG3 (clone: HP6050), IgG4 (clone: HP6025), IgM (clone: SA-DA4), IgA1 (clone: B3506B4) or IgA2 (clone: A9604D2) (all Southern Biotech, AL, USA), respectively. For the FcγR binding, a respective PE–streptavidin (Agilent Technologies) coupled recombinant and biotinylated human FcγR protein was used as a secondary probe. Excessive secondary reagent was washed away after 1h incubation, and the relative antigen-specific antibody levels were determined on an iQue analyzer (Intellicyt).

### Antibody-Dependent Complement Deposition (ADCD)

Complement deposition was performed as described before(*46*). In brief, biotinylated antigens were coupled to FluoSphere NeutrAvidin beads (Thermo Fisher) and to form immune-complexes incubated with 10 μl 1:10 diluted plasma samples for 2h at 37°C. After non-specific antibodies were washed away, immune-complexes were incubated with guinea pig complement in GVB++ buffer (Boston BioProducts, MA, USA) for 20 min at 37°C. EDTA containing PBS (15mM) was used to stop the complement reaction and deposited C3 on beads was stained with anti-guinea pig C3-FITC antibody (MP Biomedicals, CA, USA, 1:100, polyclonal) and analyzed on an iQue analyzer (Intellicyt).

### Antibody-Dependent-Neutrophil-Phagocytosis (ADNP)

Phagocytosis score of primary human neutrophils was determined as described before(*47*). Biotinylated antigens were coupled to FluoSphere NeutrAvidin beads (Thermo Fisher) and incubated with 10 μl 1:100 diluted plasma for 2h at 37°C to form immune-complexes. Primary neutrophils were derived from Ammonium-Chloride-Potassium (ACK) buffer lysed whole blood from healthy donors and incubated with washed immune complexes for 1h at 37°C. Afterwards, neutrophils were stained for surface CD66b (Biolegend, CA, USA; 1:100, clone: G10F5) expression, fixed with 4% para-formaldehyde and analyzed on a iQue analyzer (Intellicyt).

### Antibody-Dependent-THP-1 Cell-Phagocytosis (ADCP)

THP-1 phagocytosis assay was performed as described before(*48*). In brief, biotinylated antigens were coupled to FluoSphere NeutrAvidin beads (Thermo Fisher) and incubated with 10 μl 1:100 diluted plasma for 2h at 37°C to form immune complexes. THP-1 monocytes were added to the beads, incubated for 16 h at 37°C, fixed with 4% para-formaldehyde and analyzed on a iQue analyzer (Intellicyt).

### Antibody-Dependent-NK-Activation (ADNKA)

To determine Antibody-dependent NK cell activation, MaxiSporp ELISA plates (Thermo Fisher) were coated with respective antigen for 2h at RT and then blocked with 5% BSA (Sigma-Aldrich). 50 μl 1:50 diluted plasma sample or monoclonal Abs was added to the wells and incubated overnight at 4°C. NK cells were isolated from buffy coats from healthy donors using the RosetteSep NK cell enrichment kit (STEMCELL Technologies, MA, USA) and stimulated with rhIL-15 (1ng/ml, STEMCELL Technologies) at 37°C overnight. NK cells were added to the washed ELISA plate and incubated together with anti-human CD107a (BD, 1:40, clone: H4A3), brefeldin A (Sigma-Aldrich, MO, USA), and monensin (BD) for 5 hours at 37°C. Next, cells were surface stained for CD56 (BD, 1:200, clone: B159), CD16 (BD, 1:200, clone: 3G8), and CD3 (BD, 1:800, UCHT1). After fixation and permeabilization with FIX & PERM Cell Permeabilization Kit (Thermo Fisher), cells were stained for intracellular markers MIP1β (BD, 1:50, clone: D21-1351) and IFNγ (BD, 1:17, clone: B27). NK cells were defined as CD3-CD16+CD56+ and frequencies of degranulated (CD107a+), INFγ+ and MIP1β+ NK cells determined on an iQue analyzer (Intellicyt)(*49*).

### Virus neutralization

Three-fold serial dilutions ranging from 1:12 to 1:8748 were performed for each plasma sample before adding 50–250 infectious units of pseudovirus expressing the SARS-CoV-2 reference (Wuhan/wildtype) or delta variant Spike to hACE-2 expressing HEK293 cells for 1 hour. Percentage neutralization was determined by subtracting background luminescence measured in cell control wells (cells only) from sample wells and dividing by virus control wells (virus and cells only). Pseudovirus neutralization titers (pNT50) values were calculated by taking the inverse of the 50% inhibitory concentration value for all samples with a pseudovirus neutralization value of 80% or higher at the highest concentration of serum.

### Data analysis and Statistics

Data analysis was performed using GraphPad Prism (v.9.2.0) and RStudio (v.1.3 and R v.4.0). Comparisons between the adults and children were performed using Wilcoxon-signed rank test followed by Benjamini-Hochberg (BH) correction. Multivariate classification models were built to discriminate humoral profiles between vaccination arms. Prior to analysis, all data were normalized using z-scoring. Feature selection was performed using least absolute shrinkage and selection operator (LASSO). Classification and visualization were performed using partial least square discriminant analysis (PLS-DA). Model accuracy was assessed using ten-fold cross-validation. These analyses were performed using R package “ropls” version 1.20.0 (*50*) and “glmnet” version 4.0.2(*51*). Co-correlates of LASSO selected features were calculated to find features that can equally contribute to discriminating vaccination arms. Correlations were performed using Spearman method followed by Benjamini-Hochberg correction. The co-correlate network was generated using R package “network” version 1.16.0(*52*).

---

**Supplemental Figure 1.**
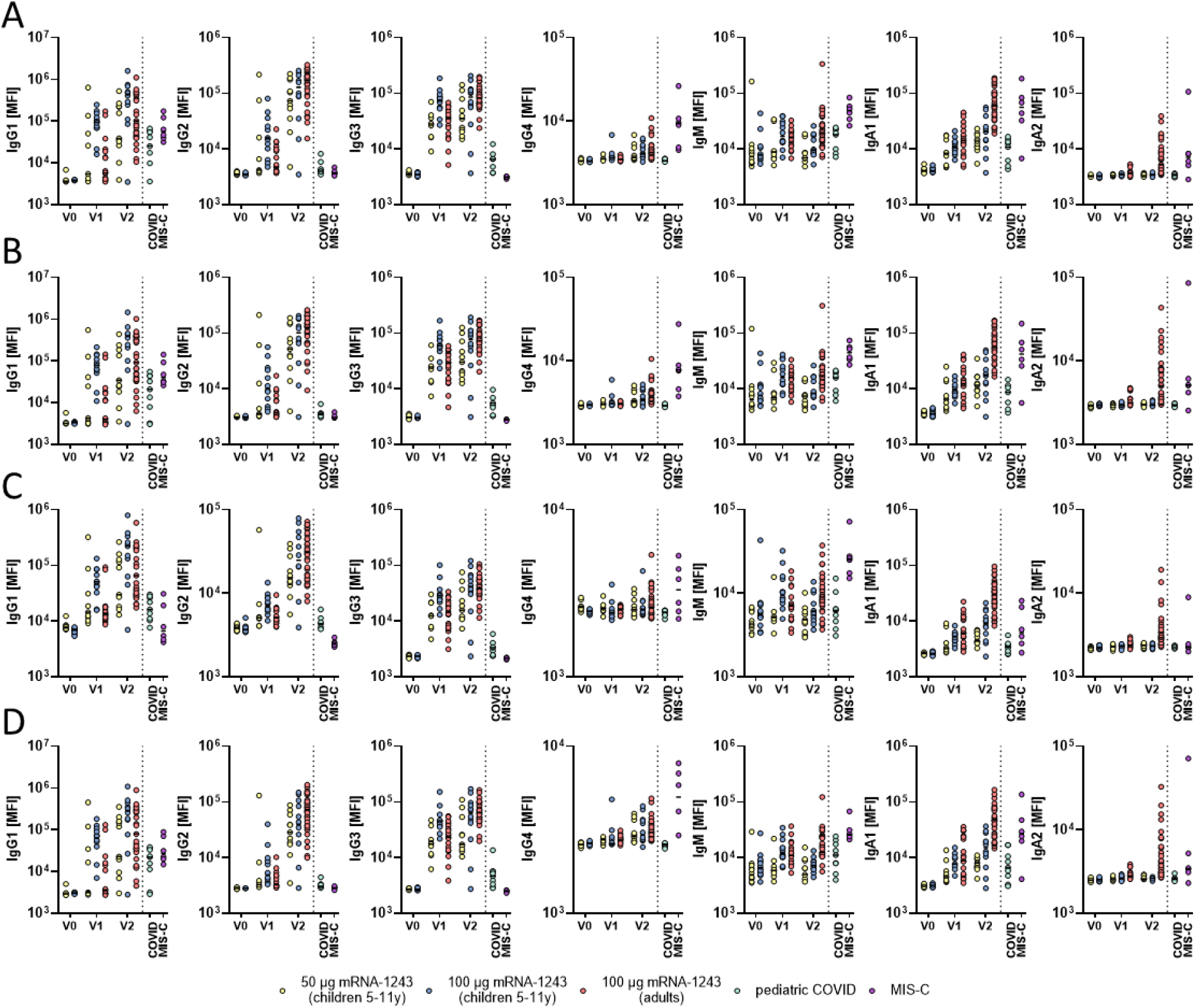
Vaccine induced antibody responses to different SARS-CoV-2 variants of concern. IgG subclass (IgG1, IgG2, IgG3, IgG4) and isotype (IgM, IgA1, IgA2) titers are shown to the receptor binding domain of wildtype (A), alpha (B), beta (C), and delta (D) variants of concern.

**Supplemental Figure 2.**
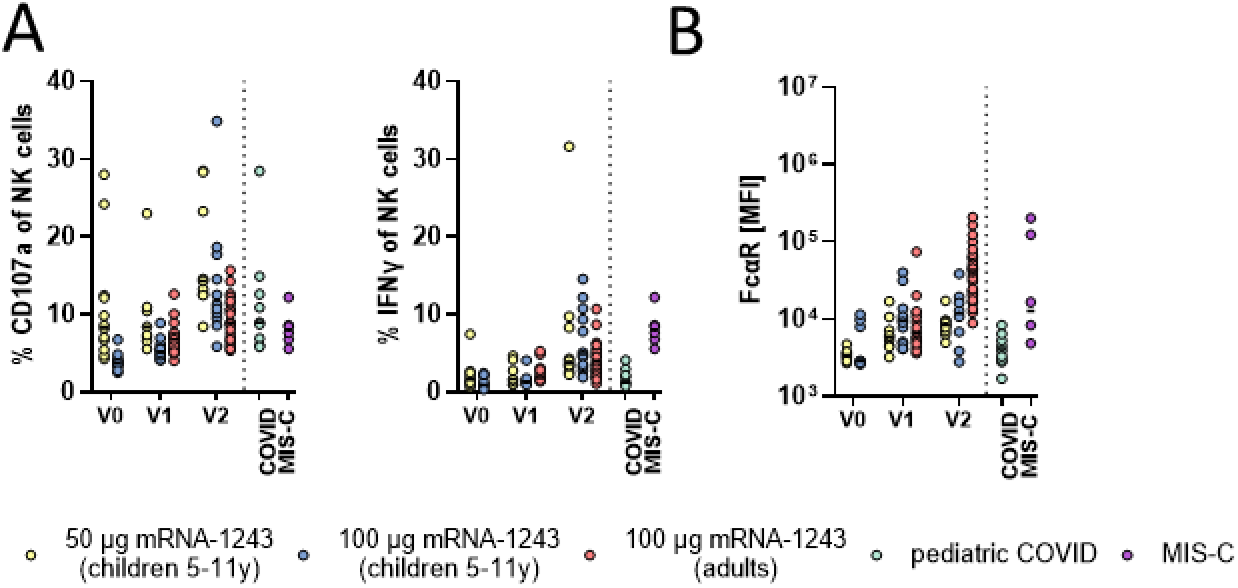
Univariate comparisons across vaccine profiles in children and adults. A) The plots show the antibody dependent NK cell activating (ADNKA, CD107a and IFN γ expression) levels in children and adults to the SARS-CoV-2 wt spike. B) The plots show the binding of SARS-CoV-2 wildtype specific IgA antibodies to FcαR by Luminex.

**Supplemental Figure 3.**
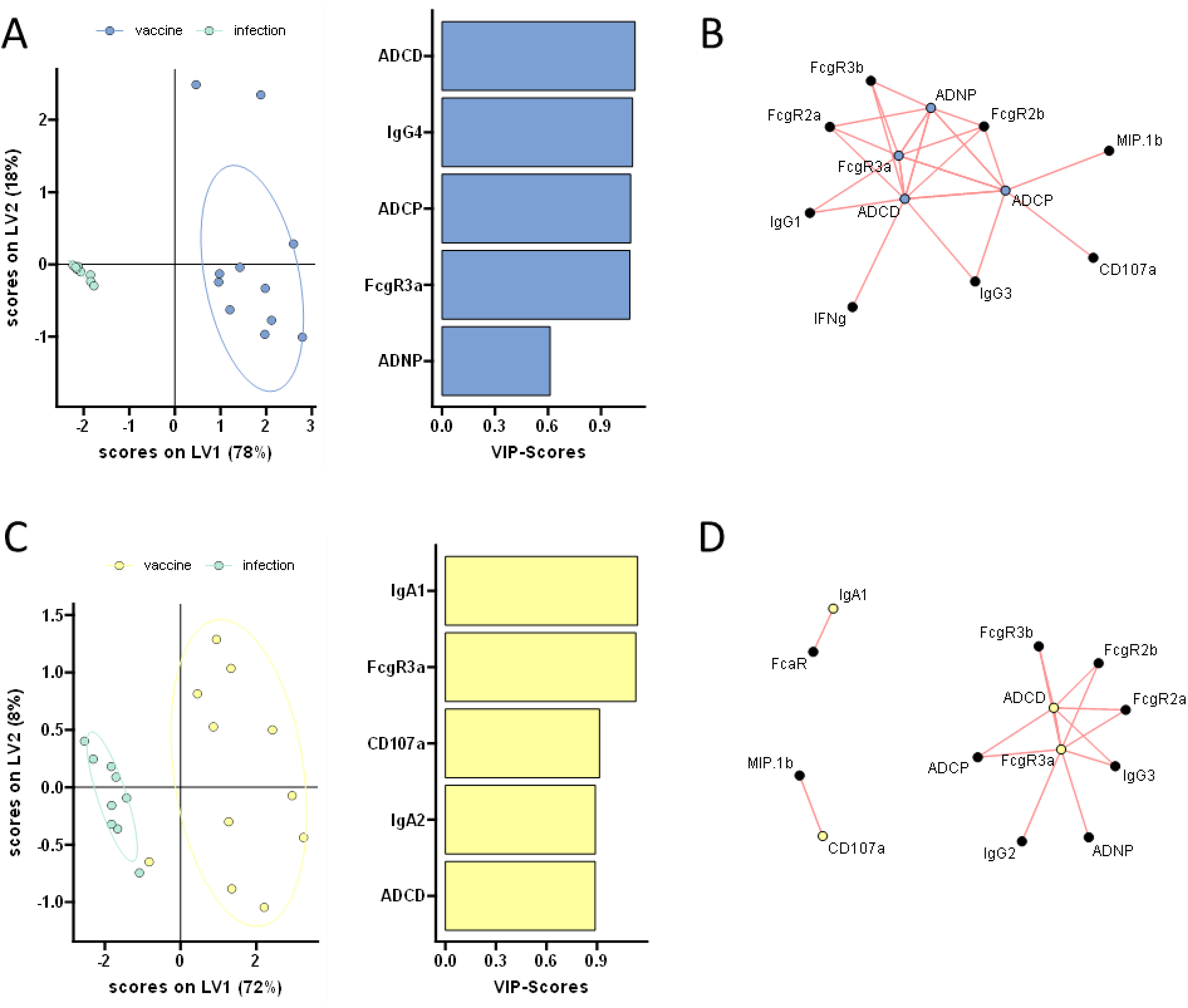
Distinct humoral profiles in naturally infected and vaccinated children. A machine learning model was built to compare SARS-CoV-2 profiles in naturally infected and 100 μg (A) or 50 μg (C) vaccinated children. A minimal set of LASSO selected SARS-CoV-2 S specific features (left panel) were first selected and used to discriminate pediatric vaccine responses (at V2) from natural infection (acute COVID) in children. Only five features were sufficient to completely separate the two groups. A co-correlate network was used to define additional features that differed in infection 100 μg (B) or 50 μg (D) vaccinated in children.

